# read_haps: using read haplotypes to detect same species contamination in DNA sequences

**DOI:** 10.1101/2020.02.11.941773

**Authors:** Hannes P. Eggertsson, Bjarni V. Halldorsson

## Abstract

**Motivation:** Data analysis is requisite on reliable data. In genetics this includes verifying that the sample is not contaminated with another, a problem ubiquitous in biology.

**Results:** In human, and other diploid species, DNA contamination from the same species can be found by the presence of three haplotypes between polymorphic SNPs. read_haps is a tool that detects sample contamination from short read whole genome sequencing data.

**Availability:** github.com/DecodeGenetics/read_haps

## 1 Introduction

Short read sequencing (SRS) is commonly used for studying human genetic variation. Reads that overlap polymorphic positions in the genome can be used to determine polymorphisms, most frequently SNPs and indels. Reads or read pairs that overlap two SNPs can be used to assign alleles to haplotypes (Halldórsson *et al.*, 2002; Lippert *et al.*, 2002). In human and other diploid species it should be possible to assign the alleles of two SNPs carried to exactly two haplotypes.

A number of reasons exist why more than two haplotypes may be observed in sequence data; Sequencing error is ubiquitous at the base pair level, reads are commonly incorrectly mapped to the reference genome, parts of the genome are variable in copy number and finally the sample may be contaminated with DNA from another sample.

Contamination is commonly detected with the tool verifyBamID Jun *et al.* (2012), which estimates contamination from the ratio of heterozygous to homozygous markers. Admixed samples generally have a high number of heterozygous markers, which may lead to overestimated contamination levels. We developed read_haps to detect contamination without relying on population genetic assumptions. Vanquish is based on principles similar to verifyBamID Jiang *et al.* (2019) and ConFindr Low *et al.* (2019) is a tool similar to read_haps trained on bacterial data.

## 2 Methodology

read_haps takes as input a sam/bam/cram file of reads aligned to a reference genome, a VCF file of variants called and a set of reliably genotyped markers. read_haps starts by finding all heterozygous markers that are reliably genotyped in the VCF file, by default conditioning the genotypes on those that have a PHRED-scaled minimum likelihood of 40 (presumed error rate at most 10^−4^) for a genotype alternate from the most likely genotype.

Only biallelic SNPs are considered and we refer to the marker alleles as 0 and 1. A two SNP haplotype can be written as *xy* where *x* is the allele at the first marker and *y* is the allele at the second marker. Two heterozygous markers can have phase 00-11 (parity) or 01-10 (non-parity), i.e. for parity phasing the individual has two haplotypes one with allele 0 on both markers and the other with allele 1 on both markers. Observing both parity and non-parity suggests the presence of three haplotypes between the two markers, but can also be explained by genotyping error and sequencing error.

To limit the impact of genotyping error we examined the sequencing data of a 15.220 individuals (Jónsson *et al.*, 2017), including 1,548 trios and determined a set of SNPs that showed low levels of inheritance errors and could generally be reliably genotyped across samples. A detailed description of this filtering is found in the Supplementary Note of (Halldorsson *et al.*, 2016) under the subheading “Description and filtering of sequencing data”.

To limit mapping and sequencing errors, only reads with high mapQ and bases with high bp quality are considered. Three haplotypes should generally not be observed in a sample from a single diploid individual, but may occur in regions of structural variation. Three haplotypes are commonly observed in a sample that contains the mixture of the DNA of two individuals. Pairs of heterozygous markers overlapped by at least two read pairs in parity and at least two reads pairs in non-parity are considered evidence for three haplotypes at the marker pair. Samples with multiple such pairs are considered contaminated, by default at most 0.2% of marker pairs can have two reads showing both parity and non-parity.

read_haps is implemented in C++ using Seqan (Döring *et al.*, 2008) and relies on htslib (Li *et al.*, 2009).

## 3 Usage

In addition to previously defined inputs, read_haps also requires a fasta file containing the reference genome, but defaults to the file genome.fa. A set of reliable markers is given along with the software. All tests are performed and tuned to Illumina WGS data, assuming –30x sequencing coverage human genome data mapped to GRCh38 using BWA mem (Li, 2013). Single sample genotyping results have been tested using GATK (McKenna *et al.*, 2010) and GraphTyper (Eggertsson *et al.*, 2017). The program has been tested with the default options for selecting which reads and variants to consider, but a number of options can also be set by the user.

A definition of the program output is given in table 1. A sample can be given the flag “FAIL” if either very few SNP pairs are read pair phasable, which generally happens due to low coverage sequencing, or due to suspected sample contamination.

**Table 1.** Definition of output from read_haps.

| column | type | explanation |
| --- | --- | --- |
| SNP_PAIRS | int | Number of read phasable SNP pairs |
| ERROR_PAIRS | int | SNP pairs with at least one read showing parity and non-parity |
| DOUBLE_ERROR_PAIR_COUNT | int | SNP pairs with at least two reads showing parity and non-parity |
| DOUBLE_ERROR_FRACTION | float | DOUBLE_ERROR_PAIR_COUNT/SNP_PAIRS |
| REL_ERROR_FRACTION | float | ERROR_PAIRS/SNP_PAIRS |
| NONSENSE_FRACTION | float | Average fraction of reads representing base pairs different from the two alleles |
| PASS_FAIL | bool | PASS/FAIL |
| REASON |  | COVERAGE or CONTAMINATION |

In cases when contamination is detected the level of contamination at each marker pair can be output using the option “–pairs”. This will point to marker pairs where there exists evidence of both parity and non-parity phasing.

## 4 Results

We ran read_haps, using the default DOUBLE_ERROR_THRESHOLD of 0.002 (0.2%), on 10,600 Icelandic samples all sequenced on Illumina NovaSeq sequencing machines. Evidence of contamination was found in 15 (0.14%) samples while 10,585 (99.86%) passed contamination checks.

We simulated (Table 2) sample contamination by creating a bam file that is a mix of bam file sequences from two individuals. The simulations were done 100 times, each time randomly selecting two files from the set of 10,600, with the contaminating sample constituting 1,2,5,10 and 20% of the total sample. Samples with 5% sample contamination or higher were always failed in our simulations, motivating our failure threshold. DOUBLE_ERROR_FRACTION was elevated at all levels in our simulations (Table 2). For comparison purposes, the results for the 10,600 samples, without contamination, are presented as contamination of 0% in Table 2.

**Table 2.**
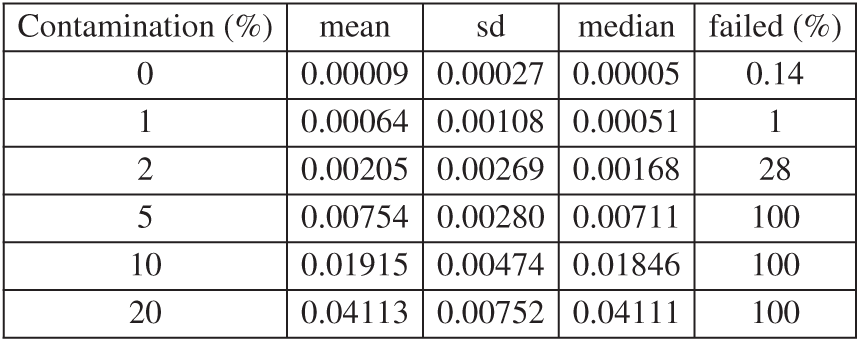
Effect of contaminating a sample on read_haps output. Reports without contamination are reported on 10,600 NovaSeq sequenced samples. Contamination is simulated at 1,2,5,10 and 20% in 100 simulations. The mean, sd and median values of DOUBLE_ERROR_FRACTION and failure rate as a function of contamination.

The FREEMIX value reported by verifyBamID (Jun *et al.*, 2012) has also been used to detect contamination. We randomly selected 1,000 of the 10,600 Icelandic samples and ran verifyBamID to estimate FREEMIX contamination levels. Mean and median FREEMIX contamination was estimated as 0.0011 and 0.0010, respectively. One sample failed at FREEMIX threshold of 2%, with FREEMIX levels of 0.02342. This sample showed an elevated double error fraction threshold 0.0014, but did not fail the default DOUBLE_ERROR_FRACTION threshold of 0.002.

We further ran verifyBamID on the 15 samples that showed evidence of contamination based in read_haps. These samples had mean and median FREEMIX contamination thresholds estimated as 0.0242 and 0.0283, respectively. 9 of these samples failed on a FREEMIX threshold of 2%, suggesting a correlation between the results of the two programs.

## Acknowledgements

We thank collaborators at deCODE genetics.

